# Functional Connectivity in Self-limited Epilepsy with Centrotemporal Spikes (SeLECTS) Increases with Epilepsy Duration and Interictal Spike Exposure

**DOI:** 10.1101/2025.07.25.666679

**Authors:** Marie E. Vasitas, Miguel S. Menchaca, Beatrice S. Goad, Christopher Lee-Messer, Zihuai He, Fiona M. Baumer

## Abstract

**Objective:** To determine the impact of epilepsy duration and interictal spikes on functional connectivity in children with Self-Limited Epilepsy with Centrotemporal Spikes (SeLECTS).

**Methods:** Connectivity was calculated from electroencephalograms (EEGs) of 68 children with SeLECTS and 65 age and sex-matched controls using the weighted phase lag index. SeLECTS EEGs were categorized by epilepsy duration (shorter or longer than 6 months) to assess progressive connectivity changes. To investigate the impact of spikes on connectivity, 19 SeLECTS patients who underwent two EEGs were analyzed longitudinally, comparing those whose spikes persisted versus resolved over time. Analyses focused on connectivity during sleep.

**Results:** Connectivity increased with epilepsy duration, being lowest in controls, intermediate in patients with shorter epilepsy duration, and highest in those with longer duration. Changes were initially greatest within the right occipital region and became more widespread with longer epilepsy duration. Longitudinally, patients with persistent spikes showed increasing connectivity over time, while those with spike resolution demonstrated decreasing connectivity, resulting in significant between-group differences.

**Conclusions:** Functional connectivity in SeLECTS increases progressively with epilepsy duration and spike exposure, suggesting that ongoing spikes drive neural network alterations.

**Significance:** Spikes are a potential treatment target to prevent progressive brain network disruption and preserve cognitive outcomes.

## 1. Introduction

Self-Limited Epilepsy with Centrotemporal Spikes (SeLECTS) is the most common focal epilepsy syndrome of childhood, comprising about 15% of pediatric epilepsy cases (Wirrell, 1998). SeLECTS is characterized by sleep-potentiated focal seizures and interictal spikes that emerge from the sensorimotor cortices, typically resolving by adolescence (Specchio et al., 2022). Despite this relatively mild clinical course, most SeLECTS patients suffer from deficits in language, attention, and reading, profoundly affecting quality of life (Baumer et al., 2018; Danielsson and Petermann, 2009; Giordani et al., 2006; Holmes, 2022; Smith et al., 2015; Teixeira and Santos, 2018). Importantly, even after seizures are controlled, cognitive impairments can still persist or even emerge (Fu et al., 2024). While seizures are infrequent, interictal spikes occur hundreds to thousands of times per night and may contribute to cognitive impairment.

Mounting evidence suggests that spikes disrupt brain networks supporting cognitive function in SeLECTS by altering both structural and functional connectivity (Besseling et al., 2013; Garcia-Ramos et al., 2015; Holmes and Lenck-Santini, 2006; Monjauze et al., 2011). Increased frontal and central connectivity correlate with cognitive deficits within the disease (Bear et al., 2019; Li et al., 2022; Ofer et al., 2018). Studies pairing functional magnetic resonance imaging (fMRI) with electroencephalogram (EEG) show increased connectivity between spike-generating motor regions and frontal regions immediately after spikes, the magnitude of which correlates with poorer language function (Vaudano et al., 2019; Xiao et al., 2016). EEG functional connectivity analyses similarly demonstrate that spikes cause immediate increases in connectivity between motor and frontal regions, and additionally show that connectivity remains elevated compared to typically-developing children even during spike-free periods of sleep (Goad et al., 2022). SeLECTS children with frequent spikes have higher connectivity compared to those with rare spikes and to typically-developing children (Dai et al., 2022). Several fMRI studies have demonstrated that functional connectivity changes with longer epilepsy duration (Wu et al., 2015; Zeng et al., 2015). Recently, an EEG study also found that connectivity progressively increased during wakefulness with longer SeLECTS duration (Garnica-Agudelo et al., 2025). However, longitudinal studies assessing connectivity over time within SeLECTS are extremely limited. While studies have established that structural connectivity progressively diverges from that in typically developing children over time (Garcia-Ramos et al., 2019, 2015; Smith et al., 2023), whether functional connectivity shows similar progression, and whether spikes drive these changes, has yet to be investigated through direct longitudinal study. Notably, prior functional connectivity studies examining the effect of epilepsy duration have focused exclusively on wakefulness, despite the fact that spikes and seizures in SeLECTS are predominantly sleep-activated.

We hypothesize that spikes drive problematic functional connectivity changes in SeLECTS, with longer exposure to spikes leading to greater connectivity deviations. To test this, we conducted a retrospective cohort study analyzing EEGs from children with SeLECTS and age/sex-matched controls without epilepsy. We employed two complementary methods: (1) Cross-sectional analysis comparing connectivity between controls and SeLECTS patients grouped by epilepsy duration (<6 months vs. >6 months since first seizure) to identify whether connectivity deviations increase with disease duration; and (2) Longitudinal analysis of SeLECTS patients with two EEGs, categorized as ‘Spikes Persist’ (spikes on both EEGs) or ‘Spikes Resolve’ (spikes on first EEG only), to evaluate whether ongoing spike activity drives divergent connectivity trajectories over time. Our analyses focused on connectivity during sleep, the time when most spikes occur and when we have previously found the most robust connectivity differences (Goad et al., 2022). We also conducted a supplementary analysis of connectivity during wakefulness, given recent evidence of progressive connectivity changes in the awake state (Garnica-Agudelo et al., 2025). We studied beta band connectivity, as it has been consistently shown to be abnormal in a wide range of SeLECTS research (Adebimpe et al., 2016, 2015; Clemens et al., 2016; Song et al., 2019), and because it can be modified with repetitive transcranial magnetic stimulation (rTMS) and thus may be a potential therapeutic target for non-invasive neuromodulation (She et al., 2025).

## 2. Methods

### 2.1 Inclusion & Exclusion Criteria

The Stanford University Institutional Review Board approved this study. Patient and control EEGs were identified using Stanford Research Repository tools. We included routine EEGs done at Lucile Packard Children’s Hospital between 2005 and 2023 monitoring children aged 3–15 years at time of EEG. Some EEGs had been included in a previous study (Goad et al., 2022), with additional EEGs identified for this study. Exclusion criteria for both groups included a history of prematurity (<35 weeks), abnormal brain MRI, other epilepsy syndromes, neurosurgery, severe brain injury, genetic disorders, profound intellectual disability, or significant medical conditions (e.g., congenital heart disease, cancer, chronic immunomodulation).

#### SeLECTS group

Eligible patients had epilepsy onset between 3–15 years of age, exhibited typical SeLECTS seizure semiology (hypersalivation, facial or hemibody twitching, or nocturnal tonic-clonic seizures), and demonstrated centrotemporal spikes on EEG. We included all initial diagnostic EEGs and follow-up EEGs that recorded both wakefulness and sleep.

#### Control group

Controls were children who underwent clinical EEGs and whom, after evaluation, had no concerns of epilepsy or serious neurologic problems (typical indications included syncope, headache, altered consciousness). Controls were also excluded if they were prescribed antiseizure medications (ASMs) or psychoactive medications.

### 2.2 EEG Acquisition

EEGs were recorded using either the Nihon-Kohden or Natus acquisition system in a standard 10-20 montage with 19 scalp electrodes at 200 or 500 Hz. All were routine clinical studies of approximately one hour or less.

### 2.3 EEG Annotation (Figure 1)

EEGs were manually annotated by trained students (BG, MV) with all annotations reviewed by a board-certified neurologist (FMB) to identify spike-free epochs in wakefulness and sleep. Spike-free epochs were defined as 500ms epochs at least 2 seconds away from any artifacts or spikes, to ensure that raised connectivity associated with spikes did not influence results. For sleep annotations, we avoided periods with vertex waves or spindles. Sleep was defined by the loss of the posterior dominant rhythm and the presence of sleep architecture (vertex waves, spindles, K-complexes). As routine EEGs were used, only stage II sleep was annotated and analyzed. To ensure representative sampling, we aimed to choose epochs at least 10 seconds apart from each other, evenly distributed across the record. Connectivity measures are sensitive to the number of epochs included (Cohen, 2014) with reasonable stabilization at 20 epochs (Ghantasala and Holmes, 2019), and thus should be matched to make valid comparisons. Since it can be challenging to find spike-free epochs in sleep in SeLECTS, we gathered 20-25 epochs of wakefulness and sleep per EEG to ensure consistency across subjects.

**Figure 1:**
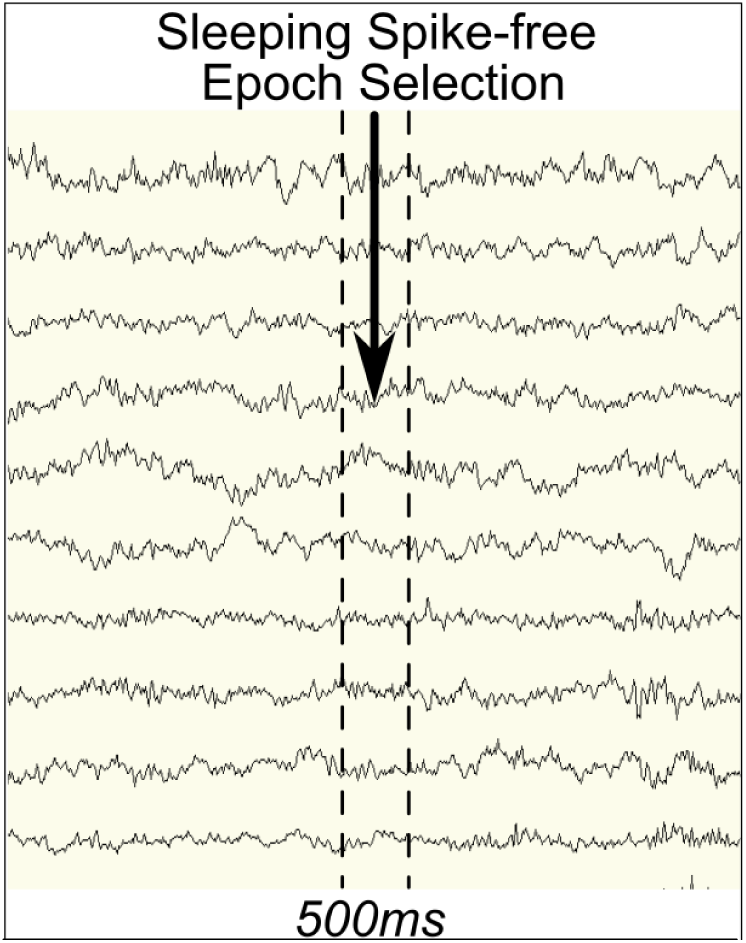
Creating EEG Epochs for Analysis. Panel shows annotation of a spike free period.

### 2.4 Clinical Factors

We recorded sex and age at EEG. For SeLECTS participants, daily ASM use at time of EEG and epilepsy duration (time since initial seizure) were also recorded.

### 2.5 Connectivity Measures

We measured connectivity using the weighted phase-lag index (wPLI), a phase based connectivity metric that is robust against the confounding effects of volume conduction in EEG (Vinck et al., 2011) and sensitive to connectivity differences in SeLECTS (Goad et al., 2022). wPLI decomposes EEG signals into the frequency band of interest, after which the phase of waveforms in each band is compared across brain regions; non-zero phase lags are considered true connectivity (Vinck et al., 2011). The beta band was chosen because it has been consistently reported as abnormal in SeLECTS (Adebimpe et al., 2016, 2015; Clemens et al., 2016; Garnica-Agudelo et al., 2025; Goad et al., 2022; Song et al., 2019) and can be reliably estimated from short epochs. EEG was band pass filtered between 12 and 30 Hz using a zero-phase, acausal filter (firwin design, Hamming window from the mne-python library (Gramfort et al., 2013) to extract the beta band, down-sampled to 200 Hz, and then the Hilbert transformation was applied (Cohen, 2014). We calculated wPLI between each of the 171 unique electrode pairs for every epoch using custom python code and then averaged across epochs (Cohen, 2014; Lee-Messer, 2021; Vinck et al., 2011). Separate connectivity values were calculated for the awake and asleep state. Data was prepared for statistical analysis using the Pandas (Mckinney, 2018) python software package facilitated by the PyArrow (Apache Arrow Team and Apache Arrow, 2022; Durant, 2022) and RAPIDS libraries (RAPIDS Development Team, 2018; Raschka et al., 2020) for quick data processing and efficient storage.

We considered connectivity at three levels of specificity, starting with each electrode-to- lectrode pair and then reducing spatial specificity by progressively averaging values, a technique that improves stability of connectivity estimates in pediatric data (Haartsen et al., 2021) and encapsulates overarching patterns of connectivity. **Pairwise connectivity** is the wPLI value between each of the 171 unique electrode pairs. **Average connectivity** considers each electrode as a node, averaging pairwise connectivity values between one electrode and the other 18 electrodes, yielding 19 values per subject. **Whole brain connectivity** is the average of all pairwise connectivity values, giving a singular connectivity value for each subject.

### 2.6 Data Analyses

In this retrospective cohort study, we conducted two complementary analyses: (1) Cross-sectional analysis comparing connectivity between controls and SeLECTS patients grouped by epilepsy duration (<6 months vs. >6 months since first seizure), including only SeLECTS EEGs with spikes present, and (2) Longitudinal analysis of SeLECTS patients who underwent two EEGs, categorized as ‘Spikes Persist’ (spikes present on both EEGs) or ‘Spikes Resolve’ (spikes present on first EEG only).

#### 2.6.1 Cross-Sectional Analysis of Epilepsy Duration

To assess if connectivity changed with disease duration, we split EEGs into 3 groups: controls, SeLECTS-Short (EEG recorded <6 months after first seizure), and SeLECTS-Long (EEG >6 months after first seizure). Prior studies have used different cut-offs for new-onset epilepsy (Ciumas et al., 2014; Garnica-Agudelo et al., 2025; Zeng et al., 2015), ranging from a few days to 12 months (Garnica-Agudelo et al., 2025). We chose a 6-month cutoff to investigate early connectivity changes with greater precision than previous studies using longer intervals, while ensuring adequate sample sizes in both groups. For the SeLECTS group, only EEGs in which spikes were recorded were included to ensure that the epilepsy had not yet been resolved. We compared controls, SeLECTS-Short, and SeLECTS-Long in pairwise, average, and whole brain connectivity, with our primary analyses focusing on connectivity during sleep and supplementary analyses focusing on connectivity during wakefulness. To account for the influence of ASM, we also conducted sensitivity analyses comparing controls to unmedicated SeLECTS patients.

#### 2.6.2 Longitudinal Analysis of Spike Persistence

To investigate whether spikes contributed to connectivity, we conducted a longitudinal analysis of children with SeLECTS who had two EEGs. All included children had spikes on their first EEG. Children who had spikes on their second EEG were categorized as “Spikes Persist” while those without spikes at second EEG were categorized as “Spikes Resolve.” Within group comparisons assessed if connectivity (pairwise, average, and whole brain) changed within each group over time, while between group comparisons assessed if the the two groups differed from each other at either of the two EEGs. Primary analyses focused on connectivity during sleep and supplementary analyses during wakefulness.

### 2.7 Statistical Analysis

Statistics were calculated using Statistical Analysis System (SAS) OnDemand for Academics (Cary, NC). Connectivity analyses were performed using generalized estimating equations (GEE) with an independent correlation matrix to account for the repeated measures of SeLECTS subjects who contributed more than one EEG to the dataset (Zeger and Liang, 1986; Zeger et al., 1988). For average and pairwise analyses, we ran separate models for each of the 19 unique electrodes and 171 unique electrode-pairs respectively and adjusted the significance threshold using Bonferroni correction (P<0.0026 for average; p<0.0003 for pairwise analyses). We compared the number of epochs included for the cross-sectional (ANOVA) and longitudinal (repeated measures ANOVA) analyses.

#### 2.7.1 Cross-sectional Analysis

We compared groups (Control, SeLECTS-Short, and SeLECTS-Long) on age, sex, and (for SeLECTS only) age of epilepsy onset and ASM use using independent t-tests, ANOVAs, and chi-square analyses. To test if connectivity changed with epilepsy duration, we fit a model with connectivity as the dependent variable and group (Control, SeLECTS-Short, and SeLECTS-Long), behavioral state (awake, asleep), and the group by behavioral state interaction as independent variables, adjusting for age, sex, and ASM use. We included both awake and asleep connectivity data in our GEE models to enhance the robustness of estimates and stratified analyses by behavioral state (Zeger et al., 1988). Our primary outcome was group differences in connectivity during sleep, and supplementary analyses assessed whole brain and average connectivity differences during wakefulness. In a second supplementary analysis, we assessed whether whole brain connectivity was correlated with age using Spearman’s correlation.

#### 2.7.2 Longitudinal Analysis

We compared the Spikes Persist and Spikes Resolve groups on sex, age at EEG 1 and EEG 2, time between the two EEGs, ASM use at EEG 1 and EEG 2, and proportion who started or stopped an ASM between the two EEGs using independent t-test and chi-square analyses. To test if spikes were associated with connectivity change, we fit a GEE model with connectivity as the dependent variable and group (Spikes Persist, Spikes Resolve), EEG (EEG 1, EEG 2), and the group by EEG interaction as independent variables, adjusting for age, sex and ASM use. We stratified analyses by group and by EEG, assessing whether there were within-group connectivity changes between EEG 1 and EEG 2 or between-group connectivity differences at either EEG 1 or EEG 2.

## 3. Results

### 3.1 Subjects

We reviewed 1019 control charts to identify 65 control children who met inclusion and exclusion criteria; all had only one EEG. All 65 controls had wakefulness on EEG, and 41 also had sleep. We reviewed 397 charts to identify 68 children who met inclusion and exclusion criteria for SeLECTS; these children contributed a total of 91 EEGs. For the cross-sectional analysis, 14 EEGs without spikes were excluded. Of the 77 remaining EEGs, 2 captured only sleep, 14 captured only wakefulness, and 61 had both, yielding 75 awake and 63 asleep SeLECTS EEGs for analysis. For the longitudinal analysis, we included SeLECTS subjects with at least 2 EEGs showing both asleep and awake data, which yielded 19 subjects with 28 EEGs.

### 3.2 Epoch Counts

For the cross-sectional analysis, there were no group differences in awake (Controls=23±3, SeLECTS-Short=23±3, SeLECTS-Long=23±3; F(2,4)=0.09, p=0.92) and asleep (Controls=24±5, SeLECTS-Short=25±7, SeLECTS-Long=22±7; F(2,2)=1.11, p=0.47) epoch counts. For the longitudinal analysis, in sleep, there were similar mean epoch counts (PersistEEG1=22±2, PersistEEG2=22±2, ResolvesEEG1=21±2, ResolvesEEG2=21±2), no significant effect of EEG number (F(1,17)=0.94, p=0.35) or group (F(1,17)=0.00, p=0.95), and no significant interaction of the two (F(1,17)=0.00, p=0.96). During wakefulness, there was also similar epoch counts (PersistEEG1=22±3, PersistEEG2=22±2, ResolvesEEG1=21±1, ResolvesEEG2=21±2), no significant effect of EEG number (F(1,17)=0.15, p=0.70) or group (F(1,17)=0.02, p=0.88), and no significant interaction between the two (F(1,17)=0.02, p=0.88).

### 3.3 Clinical Information (Tables 1-2)

#### Cross-sectional Analysis of Epilepsy Duration

Thirty-three SeLECTS EEGs were recorded within 6 months of first seizure (SeLECTS-Short) and 30 were recorded greater than 6 months from first seizure (SeLECTS-Long). Groups were evenly balanced in sex and age at EEG; however they differed significantly in ASM use and age of epilepsy onset.

**Table 1.** Counts of subjects corresponding to different demographic groups, excluding repeat EEGs without spikes.

|  | Control<br>n=65 | SeLECTS-Short<br>n=44 | SeLECTS-Long<br>n=33 | Stat | P |
| --- | --- | --- | --- | --- | --- |
| **Sex n (%)** |  |  |  |  |  |
| **Female** | 22 (34%) | 14 (32%) | 10 (30%) | $\chi^2 = 0.14$ | 0.93 |
| **Male** | 43 (66%) | 30 (68%) | 23 (70%) |  |  |
| **Age at EEG (yrs),**<br>**mean +/- SD** | 8.80 ± 2.41 | 8.32 ± 2.04 | 9.34 ± 2.03 | F = 2.02 | 0.14 |
| **Age at Onset (yrs),**<br>**mean +/- SD** | - | 8.26 ± 2.07 | 6.44 ± 2.43 | F = 12.54 | <.01* |
| **Epilepsy Duration**<br>**(yrs), mean +/- SD** | - | 0.06 ± 0.06 | 2.90 ± 1.80 | F = 110.7 | <.01* |
| **Antiseizure**<br>**Medication Use, n (%)** | - | 4 (9%) | 18 (55%) | $\chi^2 = 19.1$ | <.01* |

Statistics represent independent t-test, ANOVA and chi-square contingency tests between groups. Significant differences are indicated by \* (p<0.05); SD=Standard Deviation; yrs=Years.

**Table 2.** Clinical information of Spikes Resolve and Spikes Persist groups.

|  |  | **Spikes Resolve**<br>**n=9** | **Spikes Persist**<br>**n=10** | **Stat** | **P** |
| --- | --- | --- | --- | --- | --- |
| **Sex** | **Female** | 4 (44%) | 4 (40%) | $\chi^2 = 0.04$ | 0.85 |
|  | **Male** | 5 (55%) | 6 (60%) |  |  |
| **Age at EEG1 (yrs),**<br>**mean +/- SD** | | 8.5 ± 2.6 | 8.1 ± 1.6 | $t = 0.41$ | 0.69 |
| **Age at EEG2 (yrs),**<br>**mean +/- SD** | | 11.2 ± 2.2 | 10.3 ± 1.7 | $t = 1.49$ | 0.34 |
| **Change in Age (yrs),**<br>**mean +/- SD** | | 2.7 ± 1.8 | 1.6 ± 1.0 | $t = -0.41$ | 0.69 |
| **ASM use EEG1, n (%)** | | 3 (33%) | 1 (10%) | $\chi^2 = 1.51$ | 0.21 |
| **ASM use EEG2, n (%)** | | 7 (77%) | 6 (60%) | $\chi^2 = 0.69$ | 0.41 |
| **Change in ASM Use,**<br>**n (%)** | | 4 (44%) | 5 (50%) | $\chi^2 = 0.06$ | 0.81 |

Statistics represent independent t-tests and chi-square contingency tests. Change in Age is time between EEG 1 and EEG 2. Change in ASM use represents the number of children who started using an ASM between EEG 1 and EEG 2; no child discontinued ASM use. ASM=antiseizure medication; SD=Standard Deviation; yrs=Years.

#### Longitudinal Analysis of Spike Persistence

Nineteen SeLECTS subjects were included, 9 in the “Spikes Resolve” Group and 10 in the “Spikes Persist” group. Groups were evenly balanced in sex, age at each EEG, change in age between EEGs, ASM use at each EEG, and change in ASM use between EEGs. In both groups, some children started and no children stopped ASM use between the two EEGs.

### 3.4 Cross-Sectional Analysis of SeLECTS Duration

#### 3.4.1 Whole Brain Connectivity (Figure 2A, Supplementary Table 1)

Whole brain connectivity was lowest in controls, higher in SeLECTS-Short, and highest in SeLECTS-Long groups. There were significant differences between Controls and both SeLECTS-Short and SeLECTS-Long, as well as between SeLECTS-Short and SeLECTS-Long.

**Figure 2.**
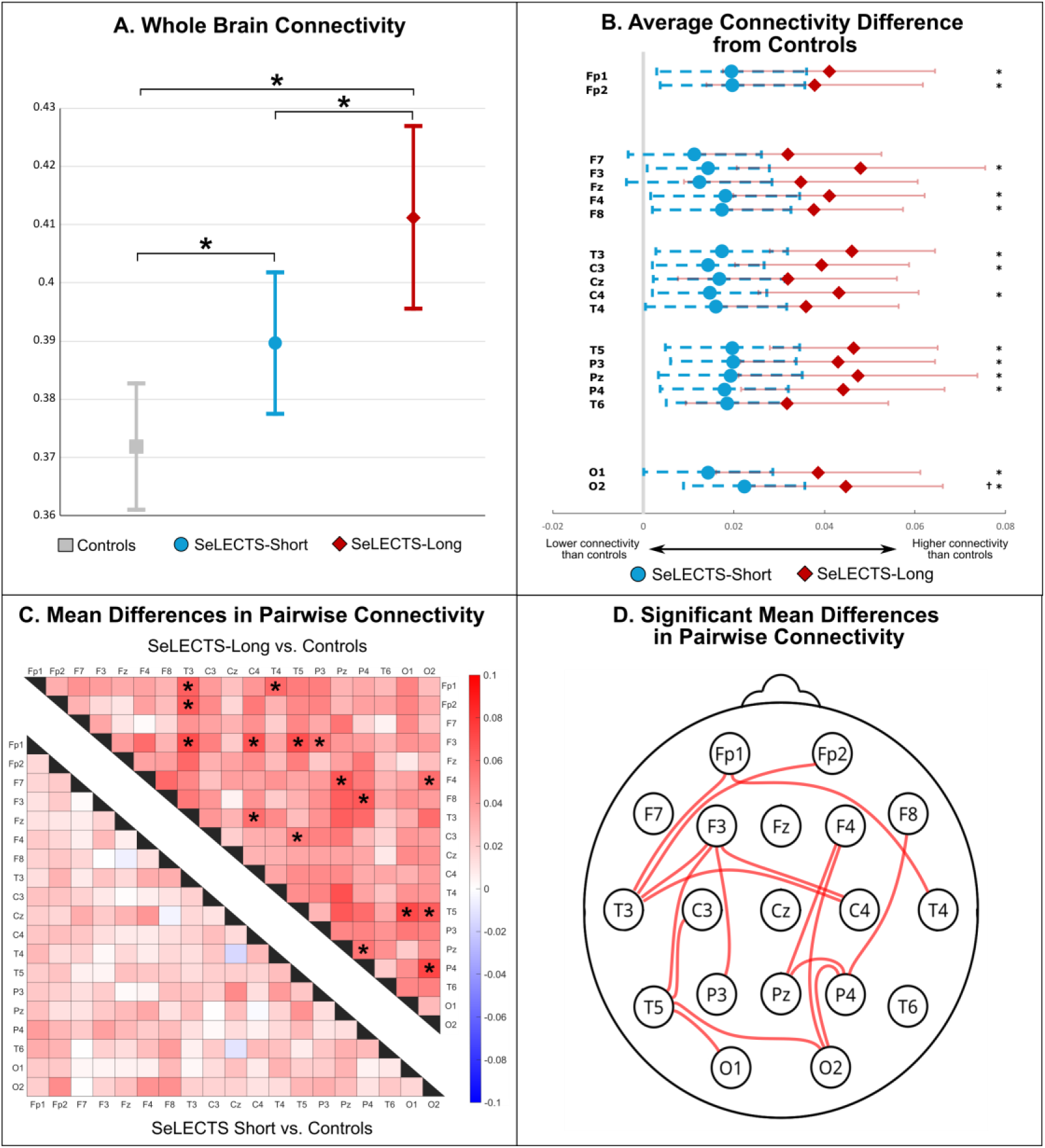
Connectivity differences between Controls, SeLECTS-Short, and SeLECTS-Long groups. Models of whole brain (**A**), average (**B)** and pairwise (**C**) connectivity, adjusted for age, sex, and antiseizure medication (ASM) use. (**A**) Least square means of whole brain connectivity and 95% confidence intervals, significant differences are denoted by a * (p <0.05). **(B)** Forest plots of group differences in average connectivity when comparing SeLECTS-Short (blue) and SeLECTS-Long (red) groups to controls. Significant differences between SeLECTS-Long and Control groups are indicated by a *, and between SeLECTS-Short and Controls by a † (p<0.0026). There are no significant connectivity differences between SeLECTS-Short and SeLECTS-Long groups. **(C)** Heat maps of group differences in pairwise connectivity. Red indicates higher connectivity in SeLECTS than controls, while blue indicates lower connectivity in SeLECTS. The diagonal divides Control comparison to SeLECTS-Long (top) and SeLECTS-Short (bottom). Significant differences are denoted by a * (p <0.0003). (**D**) Topographic map illustrating significant differences in pairwise connectivity between SeLECTS-Long and Controls. Each line represents a significant difference in pairwise connectivity between the two connected electrodes.

#### 3.4.2 Average Connectivity (Figure 2B, Supplementary Table 2)

Average connectivity was likewise lowest in controls, higher in SeLECTS-Short, and highest in SeLECTS-Long at each electrode. After multiple comparisons correction, the SeLECTS-Long group showed higher connectivity than controls at 15 of 19 electrodes. Connectivity in the SeLECTS-Short group was significantly higher than in controls at only one electrode (O2), and did not significantly differ from the SeLECTS-Long group at any electrode.

#### 3.4.3 Pairwise Connectivity (Figure 2C,D)

Pairwise connectivity was significantly higher in SeLECTS-Long compared to controls between 16 electrode-pairs. Electrodes most frequently involved in these pairs were F3, P4, T3, T5, and O2. Connectivity between the left frontal and left temporal regions (F3-T3) was significantly higher in the SeLECTS-Long vs. SeLECTS-Short group (Mean Difference=0.065, SE=0.015, p<0.0001). There were no significant pairwise differences between SeLECTS-Short and Controls.

#### 3.4.4 Awake Connectivity (Supplementary Figure 1, Supplementary Tables 3-4)

During wakefulness, whole brain connectivity was lowest within SeLECTS-Short, higher in Controls, and highest in SeLECTS-Long, although only SeLECTS-Short and SeLECTS-Long differed significantly. SeLECTS-Long average connectivity was higher than controls at three electrodes, and higher than SeLECTS-Short at seven electrodes.

#### 3.4.5 Connectivity Excluding ASM Use (Supplementary Table 5-7)

When including only SeLECTS children who did not use ASMs, whole brain connectivity was significantly higher in both SeLECTS-Short and SeLECTS-Long groups compared to controls in sleep. Average connectivity was significantly higher between SeLECTS-Short and controls at one electrode, and significantly higher between SeLECTS-Long and controls at 3 electrodes. In wakefulness, connectivity was significantly higher in SeLECTS-Long compared to controls at 6 electrodes, and significantly higher in SeLECTS-Long compared to SeLECTS-Short at 6 electrodes.

#### 3.4.6 Connectivity by Age (Supplementary Figure 2, Supplementary Table 8)

Whole brain connectivity was not correlated with age during sleep in either SeLECTS group or the control group. During wakefulness, connectivity was positively correlated with age in both Control (p=0.03) and SeLECTS-Short (p<0.001) groups, but not the SeLECTS-Long group (p=0.78).

### 3.5 Longitudinal Analysis of Spike Persistence

#### 3.5.1 Whole Brain Connectivity (Figure 3A, Supplementary Table 9)

Connectivity in the Spikes Persist group increased over time while it decreased in the Spikes Resolve group, but these within-group changes did not reach statistical significance. These divergent connectivity trajectories led to significant group differences over time, however, with the Spikes Persist group showing significantly higher connectivity than the Spikes Resolve group at EEG 2.

**Figure 3.**
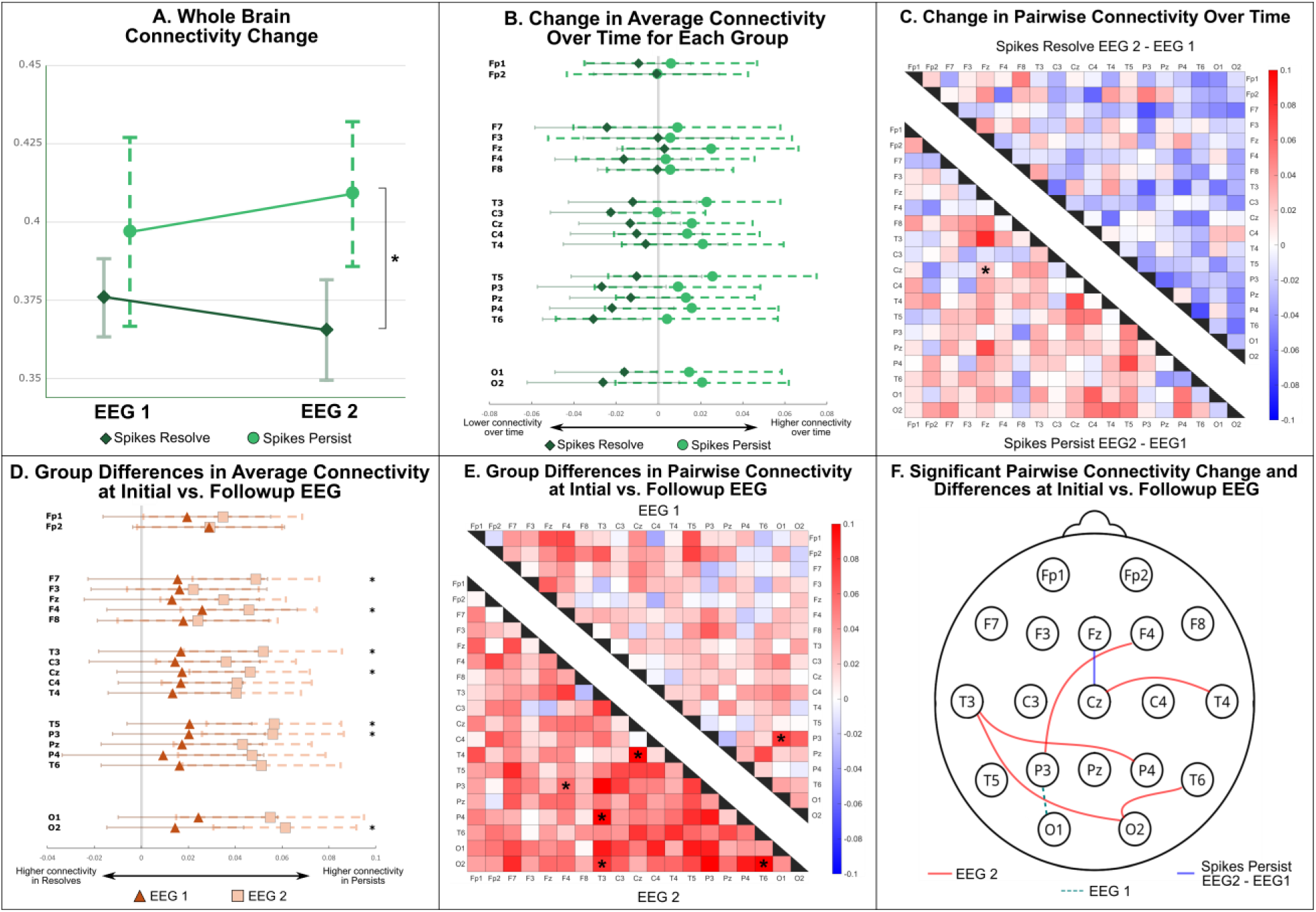
Connectivity differences over time between Spikes Resolve and Spikes Persist groups. Models of whole brain **(A)**, average **(B, D)** and pairwise **(C, E, F)** connectivity, adjusted for age, sex, and antiseizure medication (ASM) use. **(A)** Least square means of whole brain connectivity and error bars represent 95% confidence intervals, significant differences are denoted by a * (p <0.05). **(B)** Average connectivity change within groups (EEG2 - EEG1), with 95% confidence intervals. There were no significant changes. **(C)** Pairwise connectivity changes within groups (EEG2-EEG1). Blue represents a reduction in connectivity between EEGs, while red represents an increase. Significant differences are denoted by a *(p<0.0003). **(D)** Average connectivity differences between groups at EEG1 and EEG2. Negative values represent higher connectivity in the Spikes Resolve group, while a positive represents higher connectivity in the Spikes Persist Group. Significant differences at EEG2 are denoted by a * (p<0.0026). There were no significant differences at EEG1. **(E)** Pairwise connectivity differences between groups at EEG 1 and EEG 2. Red represents higher connectivity in the Spikes Persist group, while blue represents higher connectivity in Spikes Resolve. Significant differences are denoted by a * (p <0.0003). **(F)** Topographic map showing significant pairwise connectivity changes. Red and dotted red lines represent between group differences at EEG 1 and EEG2, while the solid blue line represents significant change within the Spikes Persist group between EEGs.

#### 3.5.2 Average Connectivity (Figure 3B, 3D, Supplementary Table 10)

Connectivity trended up across all electrodes in the Spikes Persist group and down in all but two of the electrodes in the between Spikes Resolve group over time, though differences were not significant. There were no significant connectivity differences between Spikes Resolve and Spikes Persist groups at EEG 1, but by EEG 2, the Spikes Persist Group showed higher connectivity at seven electrodes.

#### 3.5.4 Pairwise Connectivity (Figure 3C, 3E, 3F)

Pairwise connectivity increased between the Cz and Fz electrodes over time within the Spikes Persist group, with no significant changes in the Spikes Resolve group. A general pattern of connectivity decrease in the Spikes Resolve group and connectivity increase in the Spikes Persist group was again noted. Left parietal to occipital (P3-O1) connectivity was significantly higher in the Spikes Persist group than the Spikes Resolve group at EEG 1, whereas connectivity was significantly higher in five electrode pairs by EEG 2 (F4-P3, T3-P4, T3-O2, T4-Cz, and T6-O2).

#### 3.5.5 Awake Connectivity (Supplementary Figure 3, Supplementary Table 11-12)

When comparing whole brain connectivity during wakefulness, there were no significant differences within or between Spikes Persist and Spikes Resolve groups, although generally Spikes Persist connectivity was higher. There were no significant differences within or between groups in average connectivity during wakefulness.

## 4. Discussion

This study provides compelling evidence that functional brain connectivity in children with SeLECTS increases progressively with disease duration, and suggests that spikes drive divergent connectivity trajectories over time. Using both a cross-sectional and longitudinal approach, we demonstrate that children with longer epilepsy duration have greater connectivity compared to controls, and that connectivity changes likely depend on whether or not spikes persist. These findings support the hypothesis that interictal spikes, traditionally viewed as benign markers of SeLECTS, are a driving force behind the pathological network changes that underlie the cognitive difficulties within the disease.

### 4.1 Connectivity increases with greater duration of SeLECTS

Our cross-sectional analyses reveal a clear dose-dependent relationship between epilepsy duration and functional connectivity, with connectivity progressively increasing with longer disease duration. Studies using various imaging and neurophysiologic tools have also found progressive changes in connectivity with longer SeLECTS duration, although these changes vary by brain region and modality of study (Smith et al., 2023; Song et al., 2019; Zeng et al., 2015). In line with our results, a prior EEG study also found that beta-band connectivity progressively increases with disease duration during wakefulness (Garnica-Agudelo et al., 2025), though connectivity reductions were seen in the theta frequency band. Our study redemonstrates progressive increases in connectivity during wakefulness and additionally highlights much stronger connectivity changes during sleep. Notably, in SeLECTS of short duration, we see consistent (though mostly non-significant) elevations in sleep connectivity but unchanged or slightly reduced awake connectivity. In children with SeLECTS of longer duration, connectivity is increased in both states, and increases in sleep are of higher magnitude than those in wakefulness. The fact that both earlier and greater changes are noted in sleep compared to wakefulness may be connected to the thalamus — a critical relay station for sleep-wake cycling — which becomes progressively more connected to the cortex over time in children with SeLECTS compared to controls (Kwon et al., 2022).

In contrast to EEG studies, fMRI connectivity studies have more variable findings, and have concentrated exclusively on wakefulness. One study (Zeng et al., 2015) found regional variability in connectivity changes, with progressive increases in left frontal language connectivity, progressive decreases in default mode network, cerebellar, and occipital connectivity, and a variable pattern (initially elevated, then reduced) in the sensorimotor cortices. By contrast, a second fMRI study (Wu et al., 2015) found decreased connectivity between the motor cortex and the left inferior frontal gyrus with longer disease duration. Furthermore, *structural* connectivity, as measured by diffusion tensor imaging, appears to reliably decrease with longer disease duration (Ciumas et al., 2014; Thorn et al., 2020). Our work provides additional evidence that epilepsy disrupts typical developmental trajectories of connectivity (Ibrahim et al., 2014) and demonstrates that this effect occurs even in SeLECTS, a condition whose perception as benign has historically discouraged longitudinal investigation.

One important consideration is that children with longer disease duration also had significantly earlier epilepsy onset. Earlier onset SeLECTS may represent a more severe phenotype with greater propensity for persistent spikes and progressive connectivity changes. Furthermore, children with earlier onset experience spike exposure during more vulnerable developmental windows when brain networks are actively forming. This aligns with prior studies showing that earlier onset epilepsy is associated with worse cognitive outcomes (Garcia-Ramos et al., 2015; Hermann et al., 2008).

### 4.2 Connectivity changes evolve from focal to global with increasing duration of SeLECTS

To our surprise, we find that the most robust connectivity differences are initially seen in the right occipital region and then grow over time to involve broader networks. The recent EEG-connectivity study (Garnica-Agudelo et al., 2025) also found this pattern of spatial spread, with early increases in right parieto-occipital and the left parieto-temporal connectivity followed by later increases in the left frontal-temporal regions. The early occipital changes were unexpected, as we anticipated initial alterations would localize to the centrotemporal regions where spikes originate. These findings may suggest that the occipital cortex is more susceptible to early network disruptions in epilepsy. In support of this, prior fMRI studies have also found increased brain activity in the occipital (calcarine) cortices in children with SeLECTS (Wu et al., 2015). Alternatively, this may represent deviation from an expected developmental trajectory; typically-developing children show large age-related changes in occipitotemporal (visual network) connectivity, whereas children with SeLECTS have a flatter trajectory with increasing age (Zhang et al., 2023). Children with newly diagnosed SeLECTS have also been shown to display localized disruption of cognitive networks, predominantly in the frontal and posterior cingulate cortices, that later become widespread (Li et al., 2022). Our findings suggest that while initial connectivity alterations may not occur directly at spike foci, they rapidly propagate to distributed networks, consistent with SeLECTS being a network disorder rather than purely focal pathology.

### 4.3 Hyperconnectivity is associated with persistence of spikes

Spikes are associated with large transient connectivity increases in children with SeLECTS (Ghantasala and Holmes, 2019; Goad et al., 2022), but whether spikes specifically drive network changes has remained unclear. Our longitudinal analyses show that hyperconnectivity increases as spikes persist, and decreases once spikes resolve, providing compelling evidence for a mechanistic link between spike activity and network reorganization. Two hypotheses could explain our findings: (1) cumulative exposure to spikes could progressively drive hyperconnectivity via activity-dependent plasticity mechanisms (Knowles et al., 2022); or (2) high connectivity could represent an inherently permissive brain state that facilitates spike generation, with connectivity normalization a prerequisite for spike (and epilepsy) resolution. Our longitudinal data, which shows that connectivity decreases as spikes resolve and increases as spikes persist, is more consistent with the permissive state model. A recent fMRI study also suggests reversible network states enable spike generation, demonstrating that increased thalamocortical functional connectivity between the ventrolateral thalamus and inferior motor cortex is present in children with SeLECTS only when epilepsy is active (defined by recent seizure activity), normalizing after seizure resolution (Kwon et al., 2022). In contrast, our cross-sectional data supports the spike-driven model, as we find that connectivity increases with disease duration while generally seizure and spike frequency decrease with age (Arhan et al., 2018). Several other studies also find persistent changes in structural (Thorn et al., 2020) and functional connectivity (Chinappen et al., 2023) after spike resolution, suggesting that permanent changes have occurred. It is possible that both of these explanations are at play, with spikes creating initially reversible functional alterations that over time lead to persistent structural and functional changes. This transition may represent a critical threshold where reversible network states become entrenched developmental alterations, with important implications for the timing and targets of therapeutic intervention.

### 4.4 Clinical Implications

Our findings provide a neurobiological rationale for using spikes as a treatment target in SeLECTS to offer potential cognitive benefits. The evidence linking spikes, connectivity, and cognition operates through a clear progression: spikes cause acute cognitive disruption (Holmes and Lenck-Santini, 2006) and immediate increases in connectivity between epileptic and cognitive networks (Goad et al., 2022), with stronger spike-induced connectivity correlating with worse language function (Ibrahim et al., 2014; Xiao et al., 2016). Our study adds the critical finding that connectivity changes progress with epilepsy duration, specifically in association with persistent spike exposure, creating network alterations even during spike-free periods. The dose-dependent relationship between spikes and connectivity supports a causal role for spikes in inducing network abnormalities in SeLECTS. While the cognitive consequences of these network disturbances remain incompletely understood, evidence suggests chronic connectivity changes contribute to lasting cognitive deficits. In children with SeLECTS, disrupted connectivity correlates with poorer language function (Besseling et al., 2013), baseline network measures of connectivity predict cognitive outcomes years later (Garcia-Ramos et al., 2019), and children with persistent spikes show continued cognitive impairments even after seizure control (Baglietto et al., 2001; Fu et al., 2024). A key clinical question is whether interventions targeting spikes and connectivity can prevent or reverse progressive changes and improve cognitive outcomes. Direct manipulation of spikes is possible with ASMs and non-invasive neurostimulation: levetiracetam reduces spike frequency (Kanemura et al., 2018) and modifies spike spread to other brain regions (Wu et al., 2018) with potential language benefits (Kossoff et al., 2007), while repetitive transcranial magnetic stimulation can reduce both connectivity and spike frequency (She et al., 2025). These interventional approaches represent important next steps to determining whether modifying connectivity affects cognitive function, and whether spike suppression strategies should specifically target network normalization rather than simply reducing spike frequency.

### 4.5 Limitations

Several limitations should be considered. First, our use of clinical EEGs meant that children’s activity was not standardized during recordings. Larger connectivity differences might be apparent if we standardized for activity during wakefulness (Qi et al., 2025). Second, we limited our analysis to sensor space, which has limited spatial specificity, as we did not have participant MRIs and EEGs were low-density, clinical recordings. While more nuanced differences may emerge with source space analysis, we still saw large group differences even with sensor space analysis. Third, as we relied on clinical data, our study may be susceptible to selection bias, as children with longer duration of SeLECTS or those who underwent more than one EEG may have had more severe epilepsy. Similar results in sensitivity analyses excluding children on ASMs is reassuring, as ASM use generally correlates with disease burden in pediatric epilepsy. Fourth, our analysis focused on spike presence versus absence rather than quantifying spike frequency or characteristics. While this binary approach may be oversimplified, it reflects practical limitations of clinical EEGs, which are of limited duration. We focused on studies where sleep was recorded as spikes persist in sleep longer than wakefulness in SeLECTS. Fifth, as previously described, the 6-month cutoff used for epilepsy duration, while based on prior literature and also allowing adequate sample size in both epilepsy groups, may not represent a biologically critical time point. Sixth, in the longitudinal analysis, our modest sample size limited power to detect significant within-group changes over time, despite clear trends toward divergent trajectories. The small sample also limited our ability to explore additional factors that might influence spike persistence and connectivity trajectories. Replication in larger prospective longitudinal cohorts would strengthen confidence in these results and allow for more detailed subgroup analyses. Such longitudinal studies, as well as studies focused on manipulating spikes and connectivity, will be critical to elucidate the causal relationships between the neurophysiology of epilepsy and cognitive outcomes in children.

## Conclusion

We demonstrate that functional connectivity in SeLECTS increases progressively with epilepsy duration and spike persistence, with children showing divergent trajectories based on whether spikes resolve or persist over time. This critical evidence suggests that connectivity changes are driven by ongoing spike activity, challenging the traditional model of spikes as a benign epiphenomenon. These results support the need for future studies assessing the cognitive impact of targeted interventions — including pharmacologic and/or non-invasive neuromodulation — that manipulate spikes and connectivity. This approach would fundamentally shift our therapeutic focus from seizure control alone to actively preventing brain network disruptions, preserving long-term cognitive outcomes and enhancing quality of life in children with epilepsy.

## Acknowledgements

FMB receives funding for her research efforts from the NINDS K23NS116110, the Doris Duke Charitable Foundation, the Rita Allen Foundation, and the O’Farrel-Principe family. MV receives funding for her research from the Stanford Medscholars Research Program. CLM has received support from the Wu Tsai Neuroscience Institute and LVIS Inc, but these do not conflict with the current study.

## References

Adebimpe A, Aarabi A, Bourel-Ponchel E, Mahmoudzadeh M, Wallois F. EEG Resting State Functional Connectivity Analysis in Children with Benign Epilepsy with Centrotemporal Spikes. Front Neurosci 2016;10:143. 10.3389/fnins.2016.00143.

Adebimpe A, Aarabi A, Bourel-Ponchel E, Mahmoudzadeh M, Wallois F. Functional Brain Dysfunction in Patients with Benign Childhood Epilepsy as Revealed by Graph Theory. PLoS One 2015;10:e0139228. 10.1371/journal.pone.0139228.

Apache Arrow Team, Apache Arrow. PyArrow python bindings to the Apache Arrow project, a development platform for in-memory analytics. 2022.

Arhan E, Serdaroglu A, Ozturk Z, Aydın K, Hırfanoglu T. Serial changes in the paroxysmal discharges in rolandic epilepsy may predict seizure recurrence: A retrospective 3-year follow-up study. Epilepsy Behav 2018;82:150–4. 10.1016/j.yebeh.2018.03.014.

Baglietto MG, Battaglia FM, Nobili L, Tortorelli S, De Negri E, Calevo MG, et al. Neuropsychological disorders related to interictal epileptic discharges during sleep in benign epilepsy of childhood with centrotemporal or Rolandic spikes. Dev Med Child Neurol 2001;43:407–12. 10.1017/s0012162201000755.

Baumer FM, Cardon AL, Porter BE. Language Dysfunction in Pediatric Epilepsy. The Journal of Pediatrics 2018;194:13–21. 10.1016/j.jpeds.2017.10.031.

Bear JJ, Chapman KE, Tregellas JR. The epileptic network and cognition: What functional connectivity is teaching us about the childhood epilepsies. Epilepsia 2019;60:1491–507. 10.1111/epi.16098.

Besseling RMH, Jansen JFA, Overvliet GM, Van Der Kruijs SJM, Vles JSH, Ebus SCM, et al. Reduced functional integration of the sensorimotor and language network in rolandic epilepsy. NeuroImage: Clinical 2013;2:239–46. 10.1016/j.nicl.2013.01.004.

Chinappen DM, Ostrowski LM, Spencer ER, Kwon H, Kramer MA, Hämäläinen MS, et al. Decreased thalamocortical connectivity in resolved Rolandic epilepsy. Clin Neurophysiol 2023;153:21–7. 10.1016/j.clinph.2023.05.013.

Ciumas C, Saignavongs M, Ilski F, Herbillon V, Laurent A, Lothe A, et al. White matter development in children with benign childhood epilepsy with centro-temporal spikes. Brain 2014;137:1095–106. 10.1093/brain/awu039.

Clemens B, Puskás S, Spisák T, Lajtos I, Opposits G, Besenyei M, et al. Increased resting-state EEG functional connectivity in benign childhood epilepsy with centro-temporal spikes. Seizure 2016;35:50–5. 10.1016/j.seizure.2016.01.001.

Cohen MX. Analyzing neural time series data: Theory and practice. Cambridge, MA: MIT Press; 2014.

Dai X-J, Yang Y, Wang Y. Interictal epileptiform discharges changed epilepsy-related brain network architecture in BECTS. Brain Imaging Behav 2022;16:909–20. 10.1007/s11682-021-00566-w.

Danielsson J, Petermann F. Cognitive deficits in children with benign rolandic epilepsy of childhood or rolandic discharges: a study of children between 4 and 7 years of age with and without seizures compared with healthy controls. Epilepsy Behav 2009;16:646–51. 10.1016/j.yebeh.2009.08.012.

Durant M. Fastparquet. A python implementation of the parquet format. 2022.

Fu Y, Zhang J, Cao Y, Ye L, Zheng R, Li Q, et al. Recognition memory deficits detected through eye-tracking in well-controlled children with self-limited epilepsy with centrotemporal spikes. Epilepsia 2024;65:1128–40. 10.1111/epi.17902.

Garcia-Ramos C, Dabbs K, Lin JJ, Jones JE, Stafstrom CE, Hsu DA, et al. Network analysis of prospective brain development in youth with benign epilepsy with centrotemporal spikes and its relationship to cognition. Epilepsia 2019;60:1838–48. 10.1111/epi.16290.

Garcia-Ramos C, Jackson DC, Lin JJ, Dabbs K, Jones JE, Hsu DA, et al. Cognition and brain development in children with benign epilepsy with centrotemporal spikes. Epilepsia 2015;56:1615–22. 10.1111/EPI.13125.

Garnica-Agudelo D, Smith SDW, van de Velden D, Weise D, Brockmann K, Focke NK. Increase in EEG functional connectivity and power during wakefulness in self-limited epilepsy with centrotemporal spikes. Clin Neurophysiol 2025;171:107–23. 10.1016/j.clinph.2024.12.028.

Ghantasala R, Holmes GL. Benign Rolandic epilepsy: widespread increases in connectivity in a focal epilepsy syndrome. Epileptic Disorders 2019;21:567–78. 10.1684/epd.2019.1111.

Giordani B, Caveney AF, Laughrin D, Huffman JL, Berent S, Sharma U, et al. Cognition and behavior in children with benign epilepsy with centrotemporal spikes (BECTS). Epilepsy Res 2006;70:89–94. 10.1016/j.eplepsyres.2006.02.005.

Goad BS, Lee-Messer C, He Z, Porter BE, Baumer FM. Connectivity increases during spikes and spike-free periods in self-limited epilepsy with centrotemporal spikes. Clinical Neurophysiology : Official Journal of the International Federation of Clinical Neurophysiology 2022. 10.1016/J.CLINPH.2022.09.015.

Gramfort A, Luessi M, Larson E, Engemann DA, Strohmeier D, Brodbeck C, et al. MEG and EEG data analysis with MNE-Python. Frontiers in Neuroscience 2013;7:267. 10.3389/fnins.2013.00267.

Haartsen R, Mason L, Braithwaite EK, Del Bianco T, Johnson MH, Jones EJH. Reliability of an automated gaze-controlled paradigm for capturing neural responses during visual and face processing in toddlerhood. Developmental Psychobiology 2021;63:e22157. 10.1002/DEV.22157.

Hermann BP, Jones JE, Sheth R, Koehn M, Becker T, Fine J, et al. Growing up with epilepsy: a two-year investigation of cognitive development in children with new onset epilepsy. Epilepsia 2008;49:1847–58. 10.1111/j.1528-1167.2008.01735.x.

Holmes GL. Interictal Spikes as an EEG Biomarker of Cognitive Impairment. J Clin Neurophysiol 2022;39:101–12. 10.1097/WNP.0000000000000728.

Holmes GL, Lenck-Santini P-PP. Role of interictal epileptiform abnormalities in cognitive impairment. Epilepsy and Behavior 2006;8:504–15. 10.1016/j.yebeh.2005.11.014.

Ibrahim GM, Cassel D, Morgan BR, Smith ML, Otsubo H, Ochi A, et al. Resilience of developing brain networks to interictal epileptiform discharges is associated with cognitive outcome. Brain 2014;137:2690–702.

Kanemura H, Sano F, Ohyama T, Aihara M. Efficacy of levetiracetam for reducing rolandic discharges in comparison with carbamazepine and valproate sodium in rolandic epilepsy. Seizure 2018;62:79–83. 10.1016/j.seizure.2018.10.002.

Knowles JK, Xu H, Soane C, Batra A, Saucedo T, Frost E, et al. Maladaptive myelination promotes generalized epilepsy progression. Nat Neurosci 2022;25:596–606. 10.1038/s41593-022-01052-2.

Kossoff EH, Los JG, Boatman DF. A pilot study transitioning children onto levetiracetam monotherapy to improve language dysfunction associated with benign rolandic epilepsy. Epilepsy Behav 2007;11:514–7. 10.1016/j.yebeh.2007.07.011.

Kwon H, Chinappen DM, Huang JF, Berja ED, Walsh KG, Shi W, et al. Transient, developmental functional and structural connectivity abnormalities in the thalamocortical motor network in Rolandic epilepsy. Neuroimage Clin 2022;35:103102. 10.1016/j.nicl.2022.103102.

Lee-Messer C. EEG Connectivity Implementation 2021. https://github.com/cleemesser/eeg-connectivity-2022.

Li Y, Chen J, Sun J, Jiang P, Xiang J, Chen Q, et al. Changes in functional connectivity in newly diagnosed self-limited epilepsy with centrotemporal spikes and cognitive impairment: An MEG study. Brain Behav 2022;12:e2830. 10.1002/brb3.2830.

Mckinney W. Python for Data Analysis, Data Wrangling with Pandas, NumPy, and IPython. Sebastopol, CA: O’Reilly Media, Inc; 2018.

Monjauze CC, Broadbent H, Boyd SG, Neville BGR, Baldeweg T. Language deficits and altered hemispheric lateralization in young people in remission from BECTS. Epilepsia 2011;52:79–84. 10.1111/j.1528-1167.2011.03105.x.

Ofer I, Jacobs J, Jaiser N, Akin B, Hennig J, Schulze-Bonhage A, et al. Cognitive and behavioral comorbidities in Rolandic epilepsy and their relation with default mode network’s functional connectivity and organization. Epilepsy & Behavior 2018;78:179–86. 10.1016/j.yebeh.2017.10.013.

Qi W, Lee K, Nix KC, Menchaca M, She X, Maria LS, et al. Abnormal Elevated Connectivity During Language Processing is Associated with Poor Cognitive Performance in Children with Self-limited Epilepsy with Centrotemporal Spikes 2025:2025.06.19.660474. 10.1101/2025.06.19.660474.

RAPIDS Development Team. RAPIDS: a Collection of Libraries for End-To-End GPU Data 2018.

Raschka S, Patterson J, Nolet C. Machine Learning in Python: Main Developments and Technology Trends in Data Science, Machine Learning, and Artificial Intelligence. Information 2020, Vol 11, Page 193 2020;11:193. 10.3390/INFO11040193.

She X, Qi W, Nix KC, Menchaca M, Cline CC, Wu W, et al. Repetitive transcranial magnetic stimulation modulates brain connectivity in children with self-limited epilepsy with centrotemporal spikes. Brain Stimul 2025;18:287–97. 10.1016/j.brs.2025.02.018.

Smith AB, Bajomo O, Pal DK. A meta-analysis of literacy and language in children with rolandic epilepsy. Developmental Medicine and Child Neurology 2015;57:1019–26. 10.1111/dmcn.12856.

Smith SDW, McGinnity CJ, Smith AB, Barker GJ, Richardson MP, Pal DK. A prospective 5-year longitudinal study detects neurocognitive and imaging correlates of seizure remission in self-limiting Rolandic epilepsy. Epilepsy Behav 2023;147:109397. 10.1016/j.yebeh.2023.109397.

Song DY, Stoyell SM, Ross EE, Ostrowski LM, Thorn EL, Stufflebeam SM, et al. Beta oscillations in the sensorimotor cortex correlate with disease and remission in benign epilepsy with centrotemporal spikes. Brain Behav 2019;9:e01237. 10.1002/brb3.1237.

Specchio N, Wirrell EC, Scheffer IE, Nabbout R, Riney K, Samia P, et al. International League Against Epilepsy classification and definition of epilepsy syndromes with onset in childhood: Position paper by the ILAE Task Force on Nosology and Definitions. Epilepsia 2022;63:1398–442. 10.1111/EPI.17241.

Teixeira J, Santos ME. Language skills in children with benign childhood epilepsy with centrotemporal spikes: A systematic review. Epilepsy & Behavior 2018;84:15–21. 10.1016/j.yebeh.2018.04.002.

Thorn EL, Ostrowski LM, Chinappen DM, Jing J, Westover MB, Stufflebeam SM, et al. Persistent abnormalities in Rolandic thalamocortical white matter circuits in childhood epilepsy with centrotemporal spikes. Epilepsia 2020;61:2500–8. 10.1111/epi.16681.

Vaudano AE, Avanzini P, Cantalupo G, Filippini M, Ruggieri A, Talami F, et al. Mapping the Effect of Interictal Epileptic Activity Density During Wakefulness on Brain Functioning in Focal Childhood Epilepsies With Centrotemporal Spikes. Front Neurol 2019;10:1316. 10.3389/fneur.2019.01316.

Vinck M, Oostenveld R, van Wingerden M, Battaglia F, Pennartz CMA. An improved index of phase-synchronization for electrophysiological data in the presence of volume-conduction, noise and sample-size bias. NeuroImage 2011;55:1548–65. 10.1016/j.neuroimage.2011.01.055.

Wirrell EC. Benign epilepsy of childhood with centrotemporal spikes. Epilepsia 1998;39 Suppl 4:S32–41. 10.1111/j.1528-1157.1998.tb05123.x.

Wu T, Lim S-N, Tsai J-J, Chuang Y-C, Huang C-W, Lin C-C, et al. A randomized, double-blind, double-dummy, multicenter trial comparing the efficacy and safety of extended- and immediate-release levetiracetam in people with partial epilepsy. Seizure 2018;62:84–90. 10.1016/j.seizure.2018.09.008.

Wu Y, Ji G-J, Zang Y-F, Liao W, Jin Z, Liu Y-L, et al. Local Activity and Causal Connectivity in Children with Benign Epilepsy with Centrotemporal Spikes. PLoS ONE 2015;10:e0134361. 10.1371/journal.pone.0134361.

Xiao F, An D, Lei D, Li L, Chen S, Wu X, et al. Real-time effects of centrotemporal spikes on cognition in rolandic epilepsy. Neurology 2016;86:544–51. 10.1212/WNL.0000000000002358.

Zeger S, Liang K. Longitudinal Data Analysis for Discrete and Continuous Outcomes. Biometrics 1986;42:121–30.

Zeger SL, Liang K-Y, Albert PS. Models for Longitudinal Data: A Generalized Estimating Equation Approach. Biometrics 1988;44:1049. 10.2307/2531734.

Zeng H, Ramos CG, Nair VA, Hu Y, Liao J, La C, et al. Regional homogeneity (ReHo) changes in new onset versus chronic benign epilepsy of childhood with centrotemporal spikes (BECTS): A resting state fMRI study. Epilepsy Research 2015;116:79–85. 10.1016/J.EPLEPSYRES.2015.06.017.

Zhang Q, Li J, He Y, Yang F, Xu Q, Larivière S, et al. Atypical functional connectivity hierarchy in Rolandic epilepsy. Commun Biol 2023;6:704. 10.1038/s42003-023-05075-8.

