## Supplementary Tables for "Functional Connectivity in Self-limited Epilepsy with Centrotemporal Spikes (SeLECTS) Increases with Epilepsy Duration and Interictal Spike Exposure"

**Supplementary Table 1.** Group Differences in Whole Brain Connectivity by Epilepsy Duration in Sleep

| Comparison | Difference | SE | P |
|---|---|---|---|
| SeLECTS-Short - Controls | 0.018 | 0.006 | 0.004* |
| SeLECTS-Long - Controls | 0.039 | 0.010 | <0.001* |
| SeLECTS-Long - SeLECTS-Short | 0.022 | 0.009 | 0.025* |

Estimated by GEE least square means. Significant values denoted by a * (p<0.05).
SE=standard error; SeLECTS=Self-limited epilepsy with centrotemporal spikes.

**Supplementary Table 2.** Estimated connectivity difference between Controls and SeLECTS-Short, SeLECTS-Long groups during sleep

| | **SeLECTS-Short - Controls** | | **SeLECTS-Long - Controls** | | **SeLECTS-Long - SeLECTS-Short** | |
|---|---|---|---|---|---|---|
| | **Mean Difference (SE)** | **P** | **Mean Difference (SE)** | **P** | **Mean Difference (SE)** | **P** |
| **Fp1** | 0.021 (0.01) | 0.02 | 0.039 (0.01) | <0.001* | 0.018 (0.01) | 0.13 |
| **Fp2** | 0.022 (0.01) | 0.01 | 0.035 (0.01) | 0.003* | 0.014 (0.01) | 0.27 |
| **F3** | 0.015 (0.01) | 0.03 | 0.043 (0.01) | <0.001* | 0.028 (0.01) | 0.02 |
| **F4** | 0.019 (0.01) | 0.02 | 0.041 (0.01) | <0.001* | 0.022 (0.01) | 0.03 |
| **F7** | 0.012 (0.01) | 0.10 | 0.030 (0.01) | 0.003* | 0.017 (0.01) | 0.13 |
| **F8** | 0.018 (0.01) | 0.02 | 0.038 (0.01) | <0.001* | 0.020 (0.01) | 0.04 |
| **Fz** | 0.013 (0.01) | 0.10 | 0.032 (0.01) | 0.005 | 0.019 (0.01) | 0.10 |
| **C3** | 0.015 (0.01) | 0.02 | 0.037 (0.01) | <0.0001* | 0.023 (0.01) | 0.02 |
| **C4** | 0.015 (0.01) | 0.02 | 0.042 (0.01) | <0.0001* | 0.027 (0.01) | 0.003 |
| **Cz** | 0.017 (0.01) | 0.02 | 0.029 (0.01) | 0.02 | 0.011 (0.01) | 0.33 |
| **P3** | 0.020 (0.01) | 0.003* | 0.043 (0.01) | <0.001* | 0.023 (0.01) | 0.04 |
| **P4** | 0.019 (0.01) | 0.009 | 0.044 (0.01) | <0.0001* | 0.025 (0.01) | 0.03 |
| **Pz** | 0.020 (0.01) | 0.01 | 0.048 (0.01) | <0.001* | 0.028 (0.01) | 0.04 |
| **T3** | 0.018 (0.01) | 0.02 | 0.048 (0.01) | <0.0001* | 0.030 (0.01) | 0.002* |
| **T4** | 0.017 (0.01) | 0.03 | 0.038 (0.01) | <0.001* | 0.021 (0.01) | 0.04 |
| **T5** | 0.020 (0.01) | 0.007 | 0.047 (0.01) | <0.0001* | 0.027 (0.01) | 0.005 |
| **T6** | 0.019 (0.01) | 0.005 | 0.029 (0.01) | 0.009 | 0.010 (0.01) | 0.37 |
| **O1** | 0.015 (0.01) | 0.04 | 0.041 (0.01) | <0.001* | 0.026 (0.01) | 0.01 |
| **O2** | 0.022 (0.01) | 0.001* | 0.043 (0.01) | <0.0001* | 0.020 (0.01) | 0.04 |

Significant changes are denoted with a * (p<0.0026). Positive values indicate higher connectivity in SeLECTS patients compared to controls, or in SeLECTS-Long compared to SeLECTS-Short. SE=standard error; SeLECTS=Self-limited epilepsy with centrotemporal spikes.

**Supplementary Table 3.** Whole brain connectivity differences between Controls, SeLECTS-Short, and SeLECTS-Long during wakefulness

| Comparison | Difference | SE | P |
|---|---|---|---|
| SeLECTS-Short - Controls | -0.007 | 0.005 | 0.205 |
| SeLECTS-Long - Controls | 0.021 | 0.011 | 0.053 |
| SeLECTS-Long - SeLECTS-Short | 0.022 | 0.01 | 0.005* |

Estimated by GEE least square means. Significant values denoted by a * ($p<0.05$).
SE=standard error; SeLECTS=Self-limited epilepsy with centrotemporal spikes.

**Supplementary Table 4.** Estimated connectivity difference between Controls and SeLECTS groups during wakefulness.

| | SeLECTS-Short - Controls | | SeLECTS-Long - Controls | | SeLECTS-Long - SeLECTS-Short | |
|---|---|---|---|---|---|---|
| | Mean Difference (SE) | P | Mean Difference (SE) | P | Mean Difference (SE) | P |
| **Fp1** | -0.002 (0.01) | 0.69 | 0.031 (0.01) | 0.004* | 0.034 (0.01) | 0.001* |
| **Fp2** | -0.001 (0.01) | 0.87 | 0.032 (0.01) | 0.004* | 0.033 (0.01) | 0.002* |
| **F3** | -0.008 (0.01) | 0.15 | 0.026 (0.01) | 0.01 | 0.034 (0.01) | <0.001* |
| **F4** | -0.013 (0.01) | 0.05 | 0.019 (0.01) | 0.13 | 0.032 (0.01) | 0.006 |
| **F7** | -0.005 (0.01) | 0.36 | 0.020 (0.01) | 0.06 | 0.026 (0.01) | 0.01 |
| **F8** | -0.003 (0.01) | 0.53 | 0.027 (0.01) | 0.007 | 0.031 (0.01) | <0.001* |
| **Fz** | -0.010 (0.01) | 0.08 | 0.024 (0.01) | 0.06 | 0.034 (0.01) | 0.003* |
| **C3** | -0.004 (0.01) | 0.51 | 0.008 (0.01) | 0.49 | 0.013 (0.01) | 0.25 |
| **C4** | -0.005 (0.01) | 0.43 | 0.011 (0.01) | 0.37 | 0.016 (0.01) | 0.13 |
| **Cz** | -0.003 (0.01) | 0.61 | 0.018 (0.01) | 0.20 | 0.021 (0.01) | 0.10 |
| **P3** | -0.015 (0.01) | 0.04 | 0.012 (0.01) | 0.30 | 0.028 (0.01) | 0.007 |
| **P4** | -0.012 (0.01) | 0.09 | 0.011 (0.01) | 0.41 | 0.023 (0.01) | 0.08 |
| **Pz** | -0.010 (0.01) | 0.13 | 0.021 (0.01) | 0.13 | 0.031 (0.01) | 0.01 |
| **T3** | 0.004 (0.01) | 0.52 | 0.029 (0.01) | 0.005 | 0.025 (0.01) | 0.008 |
| **T4** | 0.002 (0.01) | 0.69 | 0.026 (0.01) | 0.004* | 0.024 (0.01) | 0.003* |
| **T5** | -0.007 (0.01) | 0.28 | 0.034 (0.01) | 0.005 | 0.042 (0.01) | <0.001* |
| **T6** | -0.012 (0.01) | 0.06 | 0.012 (0.01) | 0.34 | 0.024 (0.01) | 0.04 |
| **O1** | -0.010 (0.01) | 0.12 | 0.018 (0.01) | 0.18 | 0.027 (0.01) | 0.02 |
| **O2** | -0.007 (0.01) | 0.21 | 0.017 (0.01) | 0.21 | 0.024 (0.01) | 0.06 |

Significant changes are denoted with a * (p<0.0026). Positive values indicate higher connectivity in SeLECTS patients compared to controls, or in SeLECTS-Long compared to SeLECTS-Short patients. SE=standard error; SeLECTS=Self-limited epilepsy with centrotemporal spikes.

**Supplementary Table 5.** Antiseizure Medication Sensitivity Analysis of Group Differences in Whole Brain Connectivity by Epilepsy Duration in Sleep

| Comparison | Difference | SE | P |
|---|---|---|---|
| SeLECTS-Short - Controls | 0.016 | 0.006 | 0.011* |
| SeLECTS-Long - Controls | 0.031 | 0.010 | 0.003* |
| SeLECTS-Long - SeLECTS-Short | 0.015 | 0.011 | 0.173 |

Estimated by GEE least square means. Significant values denoted by a * (p<0.05).
SE=standard error; SeLECTS=Self-limited epilepsy with centrotemporal spikes.

**Supplementary Table 6.** Antiseizure Medication Sensitivity Analysis of Group Differences in Average Connectivity by Epilepsy Duration in Sleep

| | SeLECTS-Short - Controls | | SeLECTS-Long - Controls | | SeLECTS-Long - SeLECTS-Short | |
|---|---|---|---|---|---|---|
| | **Mean Difference (SE)** | **P** | **Mean Difference (SE)** | **P** | **Mean Difference (SE)** | **P** |
| **Fp1** | 0.019 (0.01) | 0.03 | 0.033 (0.01) | 0.01 | 0.014 (0.01) | 0.33 |
| **Fp2** | 0.020 (0.01) | 0.03 | 0.028 (0.01) | 0.03 | 0.009 (0.01) | 0.54 |
| **F3** | 0.012 (0.01) | 0.09 | 0.042 (0.01) | 0.003 | 0.030 (0.01) | 0.04 |
| **F4** | 0.017 (0.01) | 0.05 | 0.028 (0.01) | 0.02 | 0.011 (0.01) | 0.35 |
| **F7** | 0.012 (0.01) | 0.14 | 0.023 (0.01) | 0.02 | 0.011 (0.01) | 0.36 |
| **F8** | 0.017 (0.01) | 0.03 | 0.024 (0.01) | 0.03 | 0.006 (0.01) | 0.57 |
| **Fz** | 0.010 (0.01) | 0.22 | 0.024 (0.01) | 0.06 | 0.013 (0.01) | 0.29 |
| **C3** | 0.012 (0.01) | 0.05 | 0.028 (0.01) | 0.003 | -0.001 (0.01) | 0.97 |
| **C4** | 0.012 (0.01) | 0.05 | 0.028 (0.01) | 0.003* | -0.001 (0.01) | 0.97 |
| **Cz** | 0.015 (0.01) | 0.05 | 0.025 (0.01) | 0.06 | 0.010 (0.01) | 0.46 |
| **P3** | 0.019 (0.01) | 0.01 | 0.036 (0.01) | 0.01 | 0.017 (0.01) | 0.21 |
| **P4** | 0.019 (0.01) | 0.01 | 0.033 (0.01) | 0.01 | 0.015 (0.01) | 0.26 |
| **Pz** | 0.019 (0.01) | 0.02 | 0.041 (0.01) | 0.01 | 0.022 (0.02) | 0.15 |
| **T3** | 0.015 (0.01) | 0.04 | 0.043 (0.01) | <0.001* | 0.028 (0.01) | 0.004 |
| **T4** | 0.015 (0.01) | 0.06 | 0.028 (0.01) | 0.02 | 0.013 (0.01) | 0.29 |
| **T5** | 0.019 (0.01) | 0.02 | 0.042 (0.02) | <0.001* | 0.024 (0.01) | 0.04 |
| **T6** | 0.018 (0.01) | 0.01 | 0.019 (0.01) | 0.11 | 0.002 (0.01) | 0.91 |
| **O1** | 0.014 (0.01) | 0.05 | 0.028 (0.02) | 0.04 | 0.014 (0.01) | 0.29 |
| **O2** | 0.022 (0.01) | 0.002* | 0.029 (0.02) | 0.01 | 0.007 (0.01) | 0.55 |

Estimated connectivity difference between SeLECTS-Short and SeLECTS-Long groups and controls, excluding all participants with who used anti seizure medication. All values are from the individual GEE models of each electrode. Significant changes are denoted with a * ($p<0.0026$). SE=standard error; SeLECTS=Self-limited epilepsy with centrotemporal spikes.

**Supplementary Table 7.** Antiseizure Medication Sensitivity Analysis of Group Differences in Average Connectivity by Epilepsy Duration in Wakefulness

| | SeLECTS-Short - Controls | | SeLECTS-Long - Controls | | SeLECTS-Long - SeLECTS-Short | |
|---|---|---|---|---|---|---|
| | Mean Difference (SE) | P | Mean Difference (SE) | P | Mean Difference (SE) | P |
| **Fp1** | -0.004 (0.01) | 0.54 | 0.042 (0.01) | <0.001* | 0.046 (0.01) | <0.001* |
| **Fp2** | -0.002 (0.01) | 0.74 | 0.043 (0.01) | <0.001* | 0.045 (0.01) | <0.001* |
| **F3** | -0.009 (0.01) | 0.11 | 0.035 (0.01) | 0.002* | 0.044 (0.01) | <0.001* |
| **F4** | -0.013 (0.01) | 0.05 | 0.033 (0.02) | 0.04 | 0.046 (0.02) | 0.003 |
| **F7** | -0.006 (0.01) | 0.33 | 0.028 (0.01) | 0.04 | 0.033 (0.01) | 0.01 |
| **F8** | -0.003 (0.01) | 0.64 | 0.037 (0.01) | 0.001* | 0.039 (0.01) | <0.001* |
| **Fz** | -0.010 (0.01) | 0.08 | 0.037 (0.01) | 0.01 | 0.048 (0.01) | 0.001* |
| **C3** | 0.004 (0.01) | 0.51 | 0.019 (0.02) | 0.23 | 0.022 (0.02) | 0.14 |
| **C4** | -0.004 (0.01) | 0.52 | 0.020 (0.02) | 0.18 | 0.025 (0.01) | 0.09 |
| **Cz** | -0.004 (0.01) | 0.56 | 0.028 (0.02) | 0.10 | 0.032 (0.02) | 0.05 |
| **P3** | -0.014 (0.01) | 0.05 | 0.019 (0.01) | 0.17 | 0.033 (0.01) | 0.008 |
| **P4** | -0.013 (0.01) | 0.07 | 0.023 (0.02) | 0.17 | 0.037 (0.02) | 0.03 |
| **Pz** | -0.010 (0.01) | 0.13 | 0.030 (0.02) | 0.07 | 0.040 (0.02) | 0.01 |
| **T3** | 0.005 (0.01) | 0.47 | 0.036 (0.01) | 0.002* | 0.031 (0.01) | 0.006 |
| **T4** | 0.005 (0.01) | 0.43 | 0.031 (0.01) | 0.002* | 0.027 (0.01) | 0.004 |
| **T5** | -0.008 (0.01) | 0.27 | 0.042 (0.02) | 0.005 | 0.050 (0.01) | 0.001* |
| **T6** | -0.012 (0.01) | 0.08 | 0.022 (0.01) | 0.13 | 0.034 (0.01) | 0.02 |
| **O1** | 0.010 (0.01) | 0.12 | 0.029 (0.02) | 0.07 | 0.039 (0.02) | 0.01 |
| **O2** | 0.007 (0.01) | 0.21 | 0.031 (0.02) | 0.06 | 0.038 (0.02) | 0.02 |

Positive values indicate higher connectivity in SeLECTS groups compared to controls. Analysis excludes all participants who used antiseizure medications. Significant changes are denoted with * ($p<0.0026$). SE=standard error; SeLECTS=Self-limited epilepsy with centrotemporal spikes.

**Supplementary Table 8.** Correlation of age and connectivity within Controls, SeLECTS-Short, and SeLECTS-Long groups.

| | Awake | | Asleep | |
|---|---|---|---|---|
| **Group** | **ρ** | **P** | **ρ** | **P** |
| Controls | 0.34 | 0.01* | 0.17 | 0.29 |
| SeLECTS-Short | 0.41 | 0.01* | -0.09 | 0.61 |
| SeLECTS-Long | -0.05 | 0.79 | 0.11 | 0.56 |

Significant values are denoted by a * ($p<0.05$). SE=standard error, ρ = Spearman's correlation coefficient.

**Supplementary Table 9.** Within and Between Group Differences in Whole Brain Connectivity by Spike Persistence during sleep.

| Comparison | Estimate | SE | P |
|---|---|---|---|
| PersistEEG2 - PersistEEG1 | 0.012 | 0.019 | 0.519 |
| ResolveEEG2 - ResolveEEG1 | -0.014 | 0.013 | 0.247 |
| PersistEEG1 - ResolveEEG1 | 0.018 | 0.015 | 0.238 |
| PersistEEG2 - ResolveEEG2 | 0.043 | 0.013 | 0.001* |

Estimated by GEE least square means. Significant values denoted by a * ($p<0.05$). SE=standard error.

**Supplementary Table 10.** Longitudinal Analysis of Average Connectivity Differences Between Spikes Persist and Spikes Resolve Groups During Sleep

| | Within-Group Change Over Time | | | | Between-Group Differences at each EEG | | | |
|---|---|---|---|---|---|---|---|---|
| | Persists | | Resolves | | EEG1 | | EEG2 | |
| | Mean Diff. (SE) | P | Mean Diff. (SE) | P | Mean Diff. (SE) | P | Mean Diff. (SE) | P |
| **Fp1** | 0.006 (0.02) | 0.78 | -0.009 (0.02) | 0.61 | -0.020 (0.02) | 0.44 | -0.035 (0.02) | 0.02 |
| **Fp2** | 0.000 (0.02) | 0.98 | -0.001 (0.01) | 0.94 | -0.029 (0.02) | 0.08 | -0.029 (0.02) | 0.06 |
| **F3** | 0.006 (0.02) | 0.79 | -0.000 (0.02) | 0.99 | -0.016 (0.02) | 0.40 | -0.022 (0.01) | 0.12 |
| **F4** | 0.003 (0.02) | 0.87 | -0.017 (0.02) | 0.29 | -0.026 (0.02) | 0.21 | -0.046 (0.01) | 0.002* |
| **F7** | 0.009 (0.03) | 0.74 | -0.025 (0.01) | 0.04 | -0.015 (0.02) | 0.43 | -0.049 (0.01) | <0.001* |
| **F8** | 0.006 (0.03) | 0.82 | -0.001 (0.02) | 0.97 | -0.018 (0.02) | 0.34 | -0.024 (0.02) | 0.17 |
| **Fz** | 0.025 (0.02) | 0.14 | 0.003 (0.01) | 0.85 | -0.013 (0.02) | 0.50 | -0.035 (0.01) | 0.01 |
| **C3** | -0.001 (0.02) | 0.97 | -0.023 (0.02) | 0.16 | -0.014 (0.02) | 0.44 | -0.036 (0.02) | 0.02 |
| **C4** | 0.014 (0.01) | 0.25 | -0.010 (0.01) | 0.48 | -0.017 (0.01) | 0.22 | -0.041 (0.02) | 0.01 |
| **Cz** | 0.016 (0.01) | 0.30 | -0.014 (0.01) | 0.27 | -0.017 (0.01) | 0.22 | -0.046 (0.01) | <0.001* |
| **P3** | 0.009 (0.02) | 0.67 | -0.027 (0.02) | 0.10 | -0.020 (0.02) | 0.23 | -0.056 (0.02) | 0.0003* |
| **P4** | 0.016 (0.03) | 0.59 | -0.022 (0.02) | 0.22 | -0.009 (0.02) | 0.67 | -0.047 (0.02) | 0.003 |
| **Pz** | 0.013 (0.02) | 0.55 | -0.013 (0.01) | 0.25 | -0.017 (0.02) | 0.32 | -0.043 (0.01) | 0.004 |
| **T3** | 0.023 (0.02) | 0.24 | -0.012 (0.02) | 0.54 | -0.017 (0.02) | 0.35 | -0.052 (0.02) | 0.003 |
| **T4** | 0.021 (0.02) | 0.24 | -0.006 (0.02) | 0.69 | -0.013 (0.01) | 0.34 | -0.040 (0.01) | 0.004 |
| **T5** | 0.026 (0.02) | 0.09 | -0.010 (0.01) | 0.47 | -0.021 (0.01) | 0.13 | -0.057 (0.01) | <0.001* |
| **T6** | 0.004 (0.03) | 0.87 | -0.031 (0.02) | 0.07 | -0.016 (0.02) | 0.34 | -0.051 (0.02) | 0.003 |
| **O1** | 0.014 (0.02) | 0.51 | -0.016 (0.02) | 0.28 | -0.024 (0.02) | 0.16 | -0.055 (0.02) | 0.007 |
| **O2** | 0.021 (0.02) | 0.32 | -0.026 (0.01) | 0.04 | -0.014 (0.01) | 0.33 | -0.061 (0.02) | <0.001* |

For within group differences, positive values indicate increasing connectivity over time while negative values indicate reduced connectivity over time. For Between Group differences, positive values represent higher connectivity in the Spikes Persist Group while negative values represent higher connectivity in the Spikes Resolve group. All values are from GEE models of each electrode. Significant differences are denoted by a (p<0.0026). SE=standard error.

**Supplementary Table 11.** Within and Between Group Differences in Whole Brain Connectivity by Spike Persistence during Wakefulness.

| Comparison | Estimate | SE | P |
|---|---|---|---|
| PersistEEG2 - PersistEEG1 | 0.003 | 0.014 | 0.821 |
| ResolveEEG2 - ResolveEEG1 | 0.002 | 0.010 | 0.840 |
| PersistEEG1 - ResolveEEG1 | 0.021 | 0.012 | 0.095 |
| PersistEEG2 - ResolveEEG2 | 0.022 | 0.013 | 0.090 |

Estimated by GEE least square means. Significant values denoted by a * ($p<0.05$).
SE=standard error.

**Supplementary Table 12.** Longitudinal Analysis of Average Connectivity Differences Between Spikes Persist and Spikes Resolve Groups During Wakefulness

| | Within-Group Change Over Time | | | | Between-Group Differences at each EEG | | | |
|---|---|---|---|---|---|---|---|---|
| | Persists | | Resolves | | EEG1 | | EEG2 | |
| | Mean Diff. (SE) | P | Mean Diff. (SE) | P | Mean Diff. (SE) | P | Mean Diff. (SE) | P |
| **Fp1** | -0.006 (0.02) | 0.72 | 0.007 (0.01) | 0.60 | 0.026 (0.01) | 0.07 | 0.014 (0.01) | 0.36 |
| **Fp2** | 0.011 (0.01) | 0.41 | 0.000 (0.01) | 0.98 | 0.011 (0.01) | 0.35 | 0.021 (0.02) | 0.18 |
| **F3** | 0.019 (0.02) | 0.25 | 0.019 (0.01) | 0.03 | 0.017 (0.01) | 0.13 | 0.016 (0.01) | 0.20 |
| **F4** | -0.003 (0.01) | 0.82 | -0.002 (0.01) | 0.86 | 0.021 (0.02) | 0.17 | 0.021 (0.02) | 0.19 |
| **F7** | 0.005 (0.02) | 0.76 | -0.020 (0.01) | 0.09 | 0.013 (0.01) | 0.31 | 0.038 (0.02) | 0.01 |
| **F8** | 0.011 (0.02) | 0.53 | -0.002 (0.02) | 0.92 | -0.003 (0.02) | 0.83 | 0.010 (0.02) | 0.59 |
| **Fz** | 0.017 (0.02) | 0.29 | 0.006 (0.01) | 0.56 | 0.019 (0.01) | 0.18 | 0.030 (0.02) | 0.06 |
| **C3** | -0.008 (0.01) | 0.54 | -0.006 (0.01) | 0.56 | 0.028 (0.01) | 0.04 | 0.026 (0.02) | 0.08 |
| **C4** | -0.001 (0.01) | 0.96 | 0.008 (0.01) | 0.39 | 0.029 (0.01) | 0.02 | 0.021 (0.01) | 0.09 |
| **Cz** | 0.017 (0.02) | 0.30 | 0.004 (0.02) | 0.80 | 0.010 (0.01) | 0.50 | 0.023 (0.02) | 0.21 |
| **P3** | -0.000 (0.02) | 0.99 | -0.010 (0.01) | 0.26 | 0.011 (0.01) | 0.47 | 0.021 (0.01) | 0.14 |
| **P4** | -0.009 (0.02) | 0.67 | -0.016 (0.02) | 0.35 | 0.021 (0.02) | 0.25 | 0.028 (0.02) | 0.13 |
| **Pz** | 0.018 (0.02) | 0.23 | 0.012 (0.02) | 0.47 | 0.020 (0.02) | 0.21 | 0.026 (0.02) | 0.14 |
| **T3** | 0.007 (0.02) | 0.74 | 0.009 (0.01) | 0.44 | 0.011 (0.01) | 0.32 | 0.010 (0.01) | 0.52 |
| **T4** | 0.013 (0.02) | 0.45 | 0.007 (0.01) | 0.56 | 0.006 (0.01) | 0.62 | 0.012 (0.01) | 0.42 |
| **T5** | 0.015 (0.02) | 0.45 | -0.006 (0.02) | 0.71 | 0.007 (0.02) | 0.70 | 0.028 (0.01) | 0.05 |
| **T6** | -0.008 (0.02) | 0.66 | -0.005 (0.02) | 0.74 | 0.012 (0.02) | 0.55 | 0.009 (0.01) | 0.53 |
| **O1** | -0.012 (0.02) | 0.49 | -0.012 (0.01) | 0.19 | 0.017 (0.02) | 0.27 | 0.017 (0.02) | 0.29 |
| **O2** | -0.007 (0.02) | 0.70 | -0.005 (0.02) | 0.81 | 0.015 (0.02) | 0.37 | 0.012 (0.02) | 0.57 |

For the Within-Group differences, positive values indicate increasing connectivity over time while negative values indicate reduced connectivity over time. For Between-Group differences, positive values represent higher connectivity in the Spikes Persist Group while negative values represent higher connectivity in the Spikes Resolve group. All values are from the GEE models of each electrode. Significant differences are denoted by a ($p<0.0026$). SE=standard error.
