## Supplementary Figures for "Functional Connectivity in Self-limited Epilepsy with Centrotemporal Spikes (SeLECTS) Increases with Epilepsy Duration and Interictal Spike Exposure"

**Supplementary Figure 1**: Connectivity differences in Controls, SeLECTS-Short, and SeLECTS-Long groups during wakefulness.

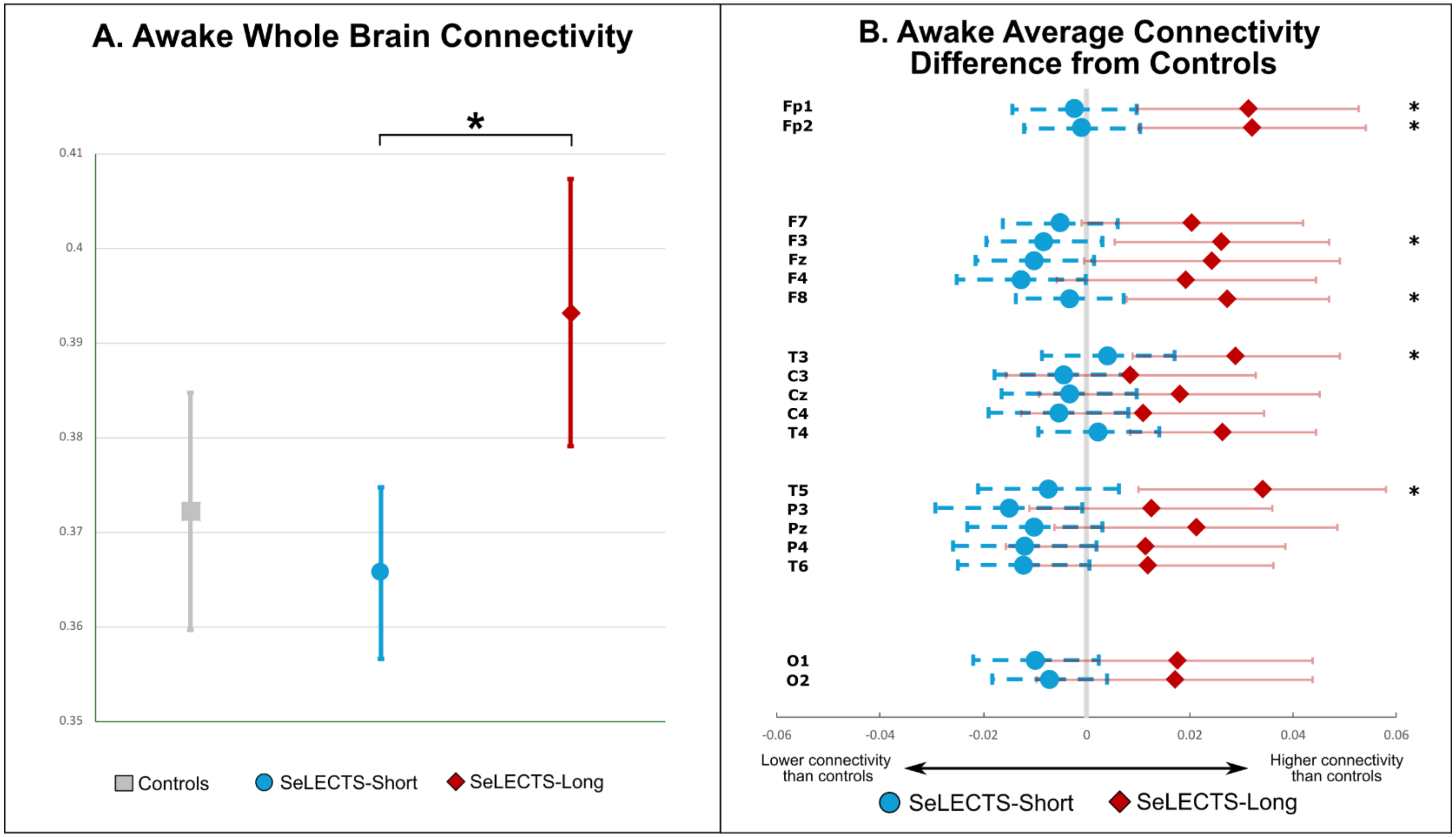


Models of whole brain (**A**) and average (**B**) connectivity during wakefulness, adjusted for age, sex, and antiseizure medication (ASM) use. (**A**) Least squares means connectivity and 95% confidence intervals. Significant differences denoted by a *(p <0.05). **(B)** Forest plots of group differences in average connectivity when comparing SeLECTS-Short (blue) and SeLECTS-Long (red) groups to controls. Significant differences between SeLECTS-Long and Control groups are indicated by a *, and between SeLECTS-Short and SeLECTS-Long by a † (p<0.0026). Note that † does not indicate a difference between SeLECTS-Short and controls, as was the case in Figure 2.

**Supplementary Figure 2.** Whole brain connectivity vs. age at EEG within SeLECTS and Controls, in both wakefulness and sleep.

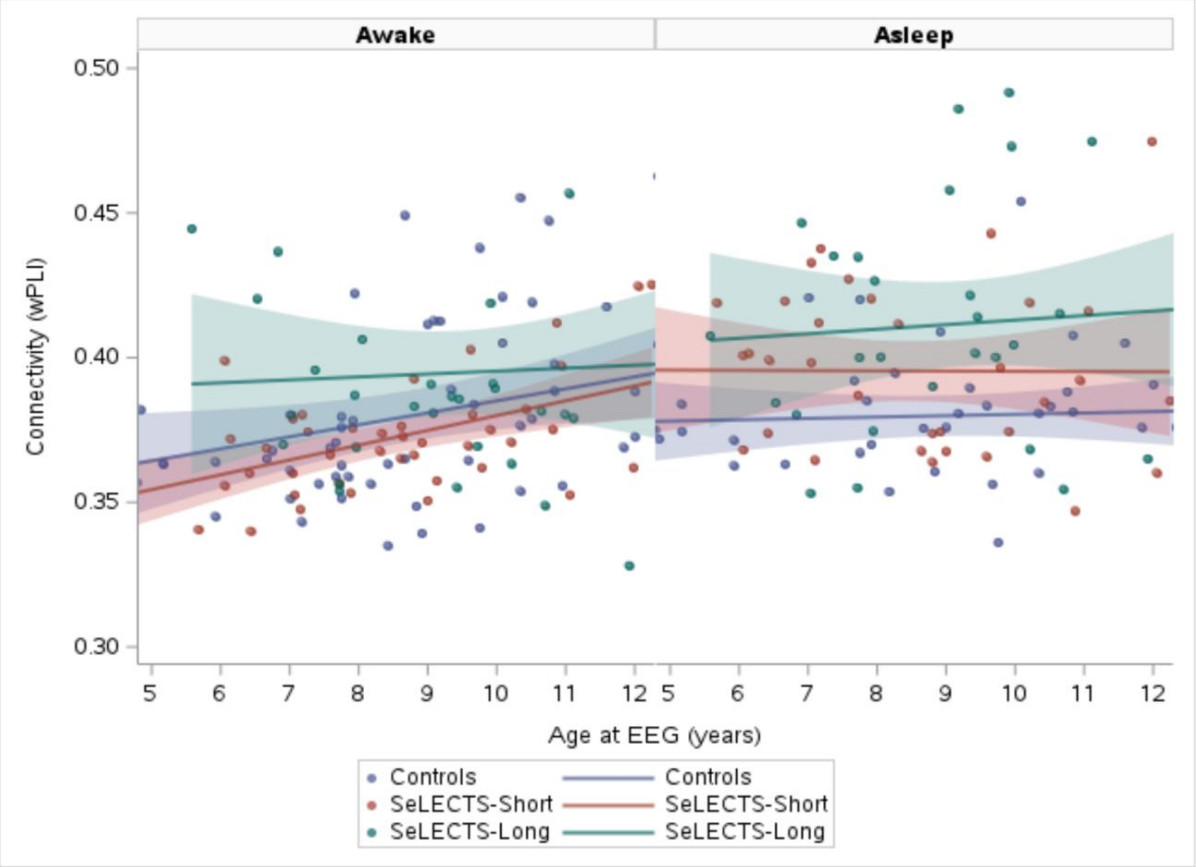


Regression lines show the relationship between age and connectivity for each group. Although linear regression lines are displayed for visualization, statistical analyses used Spearman's rank correlation (Supplementary Table 8) due to non-normal distribution of connectivity values. Controls and SeLECTS-Short had a significant correlation during wakefulness ($p < 0.01$)

**Supplementary Figure 3.** Awake Connectivity differences between Spikes Resolve and Spikes Persist groups.

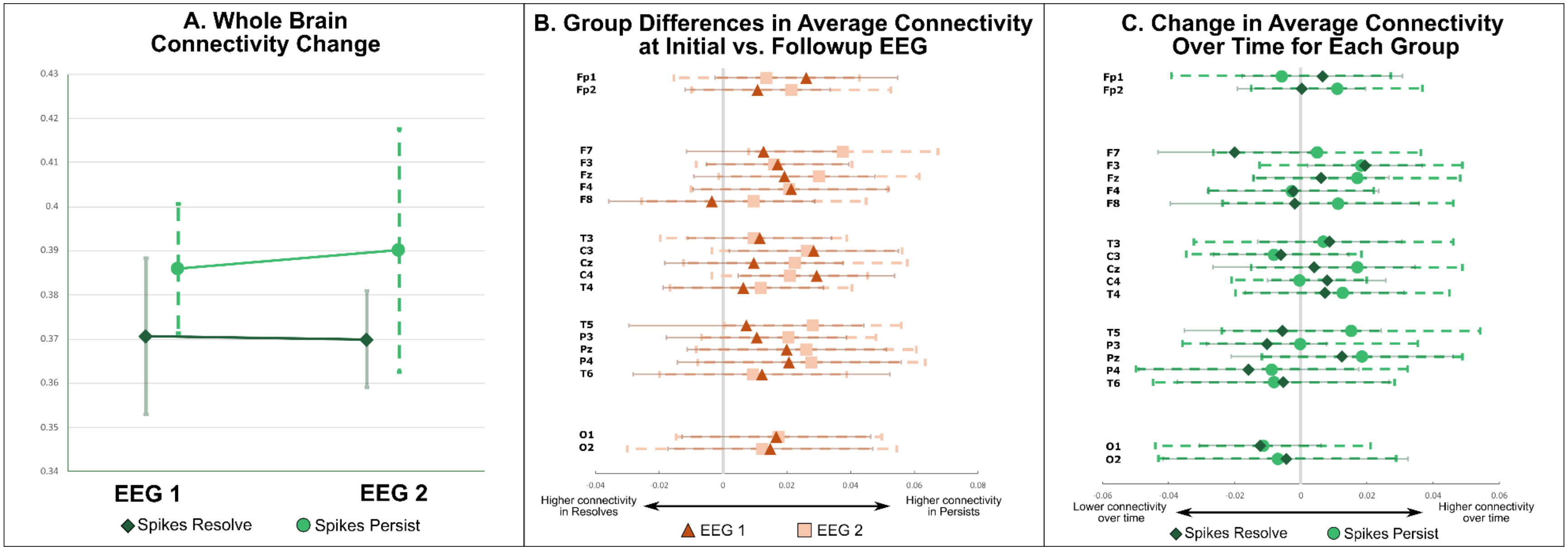


Models of whole brain **(A)** and average **(B, C)** connectivity, adjusted for age, sex, and antiseizure medication (ASM) use. **(A)** Least square means of whole brain connectivity and error bars represent 95% confidence intervals. There were no significant differences. **(B)** Average connectivity change within groups (EEG2 - EEG1), with 95% confidence intervals. There were no significant changes. **(C)** Average connectivity differences between groups at EEG1 and EEG2. Negative values represent higher connectivity in the Spikes Resolve group, while a positive represents higher connectivity in the Spikes Persist Group. There were no significant differences.
